# KANN: estimation of genetic ancestry profiles by nearest neighbor regression

**DOI:** 10.1101/2025.08.27.671485

**Authors:** Juha Riikonen, Sini Kerminen, Aki Havulinna, Matti Pirinen

## Abstract

State-of-the-art methods for inferring individual-level genetic ancestry are based on statistical models for haplotype data. Unfortunately, these methods are computationally demanding, making them impracticable for biobank-scale analyses. In this paper we describe KANN, an efficient k-nearest neighbor regression method for individual-level ancestry estimation with respect to predefined source populations using only principal components of genetic structure. Contrary to the existing tools that can only use reference samples with discrete source population assignment, KANN enables the use of reference samples with continuous ancestry profiles across multiple source populations. We illustrate KANN on a data set of 18,125 Finnish samples from THL Biobank, estimating ancestry profiles across up to 10 Finnish source populations. KANN’s ancestry estimates agree well with the haplotype-based method SOURCEFIND, showing a correlation of at least 0.859 in all 10 source populations, making KANN a promising tool for ancestry estimation in large-scale genomic studies.

## 1 Introduction

The genetic variation currently observed in human populations has been shaped by historical migrations and admixture events. Understanding and quantifying this genetic variation is essential in population genetics and genomic medicine. For example, the performance and reliability of polygenic risk scores (PRS), that have been successfully used for quantifying the genome-wide contribution to complex diseases and traits [1], depend heavily on the genetic ancestry of individuals [2, 3]. Therefore, the detailed characterization of individual-level ancestry is essential for selecting the most suitable ancestry-matched PRS model for each individual. In addition, the general public has shown great interest in their own genetic ancestry, as millions of individuals have already taken a direct-to-consumer genetic test provided by companies with affordable services and easily accessible online platforms [4]. Thus, there is a clear need to scale accurate ancestry estimation methods to biobank-scale data sets.

Various computational tools have been developed for the purpose of genetic ancestry inference. SOURCEFIND [5] and RFMix [6] are widely used methods that utilize individual-level haplotype information. Such haplotype-based methods are able to distinguish fine-scale genetic structure but suffer from slow runtime. Compared to the haplotype-level models, methods that work with individual-level genotype data (e.g. iAdmix [7]) and ADMIXTURE [8]) can improve on runtime but still they remain computationally heavy for large samples.

A recent approach called Rye [9] introduced the idea of basing genetic ancestry inference on individuals’ scores on principal components of genetic structure without a need to access variant-level data. This enabled more efficient computation, making Rye applicable to biobank-scale inference. Rye was shown to run over 50 times faster than ADMIXTURE when applied to a data set of 488,221 UK Biobank participants, while the accuracy remained comparable to the existing methods (RFMix, iAdmix and ADMIXTURE) [9].

As an input, Rye requires a set of reference individuals that are each assigned to represent a single ancestral source population. This can be a sufficient approach when only discrete population information is available and the ancestral populations are carefully defined to ensure they are relatively homogeneous and minimally admixed with one another. However, the discrete assignment likely over- simplifies the complexity of population structure we observe in modern human populations. A more flexible approach would be to use a continuous representation of ancestry, where each reference individual is characterized by an ancestry profile describing the proportion of genetic ancestry inherited from each of the source populations. This framework would allow for a more detailed depiction of genetic ancestry and has the potential to increase the precision of ancestry inference, especially in highly admixed populations.

In this work, we develop a new method for *k*-nearest neighbor regression for ancestry estimation (KANN), that works in the space of principal components of genetic structure. Importantly, KANN can utilize admixed reference individuals who have proportions of genetic ancestry inherited from multiple source populations. As a special case, KANN is applicable to the conventional ancestry format where each reference sample is assigned to exactly one population, allowing a direct comparison with Rye.

We apply KANN to a data set from THL biobank that contains fine-scale ancestry profiles with respect to 10 Finnish subpopulations made by SOURCEFIND for 18,125 individuals. We compare the results from KANN and Rye to SOURCEFIND, and we interpret the observed differences with respect to the known genetic and geographical relationships of the Finnish subpopulations.

## 2 Materials and Methods

### 2.1 Software availability

KANN is available as a software package in R-language at https://github.com/riikonenj/KANN.

### 2.2 Data

We used data from the biobank of the Finnish Institute for Health and Welfare (THL), consisting of 16,962,023 variants measured on 51,852 individuals together with the birth locations up to a municipality-level precision. The data originated from Finnish cohort studies: FINRISK (n = 30,867), GeneRISK (n = 7,255), FinHealth 2017 (n = 6,155) and Health 2000 (n = 7,575). All individuals included in this study had given a written consent.

### 2.3 PCA

#### Variants

Variant quality control was performed using PLINK 2.0 software [10]. We considered only biallelic variants found in the autosomal genome. First, we excluded variants with minor allele frequency (MAF) *<* 5%, Hardy-Weinberg p-value *<* 1e-6 and imputation information score *<* 95%. To consider only independent variants, we removed known regions of long-range linkage-disequilibrium (LD) [11]. Further, we pruned the remaining variants so that the squared pairwise correlation *r*^2^ *≤* 0.2. After filtering, 94,843 variants remained.

#### Samples

Starting from 51,852 samples, we considered only those whose municipality of birth was in Finland or in the eastern regions ceded to the Soviet Union during the Second World War. We estimated the sample heterozygosity using a method of moments F-statistic and excluded samples whose F-statistic deviated *>* 4 standard deviations (SD) from the mean. We used the KING kinship coefficient threshold *ϕ >* 0.0442 [12] to prune out the relatives starting from the third degree. After filtering, 38,113 unrelated samples remained.

#### PCA

Following the variant and sample filtering, we applied principal component analysis (PCA) on the genotypes of the 38,113 unrelated samples. The PCA was run using PLINK 2.0 software with the approximation modifier (`--pca approx`). We extracted the first 10 principal components (PCs) and the corresponding eigenvalues. An additional 9,837 non-monozygotic samples with excess kinship or homozygosity were afterwards projected on the PCs, resulting in a total of 47,950 samples with the PC values. Finally, the PCs were standardized to have zero mean and unit variance.

### 2.4 SOURCEFIND ancestry profiles

A fine-scale genetic population structure in Finland was previously determined [13] using the FINRISK study cohorts included in our data. In short, a subset of 2,741 carefully chosen reference individuals were allocated to three sets of genetically and geographically well-defined reference groups in an iterative process utilizing the ChromoPainter and FineSTRUCTURE software [14]. The reference sets characterize the fine-scale genetic ancestry in Finland using two (refset 2, n = 1,472), six (refset 6, n = 1,026) and ten (refset 10, n = 1,236) reference groups per set. SOURCEFIND [5] was further used to estimate two-, six-, and ten-dimensional ancestry profiles for a total of 18,463 samples, including the reference samples themselves. SOURCEFIND was run with 50,000 burn-in iterations, 200,000 sample iterations and results were recorded after every 5,000 iterations. When estimating ancestry for each of the reference group samples, the sample itself was left out from its reference group.

We had both our own PC coordinates and the SOURCEFIND profiles from [13] available for 18,125 FINRISK samples. Out of these samples, 1,798 were among the reference samples (1,443, 1,006 and 1,214 had a discrete population assignment with respect to 2, 6 and 10 source populations, respectively). The data quality control and pre-processing steps are described in Supplementary Fig. S1. The birth municipalities of the 1,214 samples forming the 10 source populations are depicted in Fig. 1A. Fig. 1B shows the same samples on the first two PCs. Supplementary Fig. S2 and Supplementary Fig. S3 show the same information for 2 and 6 source populations, respectively.

**Figure 1:**
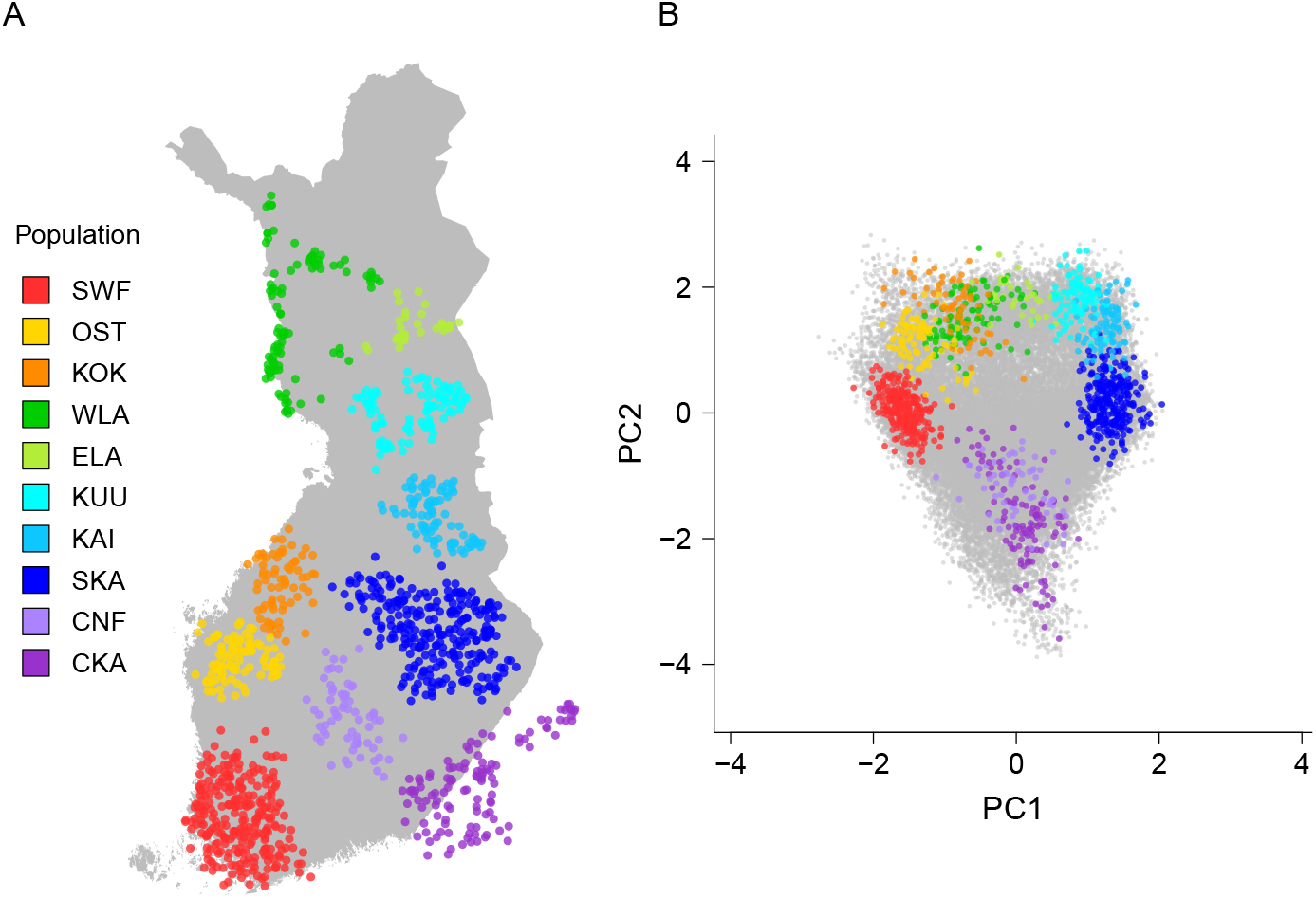
A) Map of Finland with the geographical locations of the reference samples (n = 1,214) allocated to 10 Finnish source populations. The points depict the mean coordinates of the parents’ municipalities of birth. B) The same individuals highlighted on the first two principal components. The samples not included in the reference groups are depicted in grey colour. SWF: Southwestern Finland, OST: Ostrobothnia, KOK: Kokkola, WLA: West Lapland, ELA: East Lapland, KUU: Kuusamo, KAI: Kainuu, SKA: Savo-Karelia, CNF: Central Finland, CKA: Ceded Karelia.

### 2.5 KANN algorithm

#### 2.5.1 k-nearest neighbor regression

Ancestry estimation with KANN leverages the known ancestry profiles of *n*_*R*_ reference samples, each having a proportion of their genetic ancestry assigned to *S* different source populations. We represent the ancestry profile of reference sample *j ≤ n*_*R*_ as

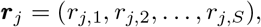

where *r*_*j,s*_ describes the ancestry proportion of sample *j* with respect to the source population *s ≤ S*. Leveraging the known ancestry information of the reference samples, the aim is to estimate, for each query sample *i ≤ n*_*Q*_, a corresponding ancestry profile:

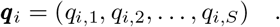

To apply k-nearest neighbor regression for ancestry estimation, we define the distance *d*(*i, j*) *≥* 0 between query sample *i* and reference sample *j* as their Euclidean distance in the *M* -dimensional PC-space, where the distance along each component *m* is weighted with the component’s eigenvalue *λ*_*m*_:

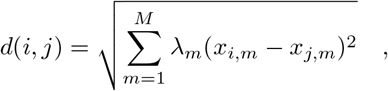

where *x*_*i,m*_ and *x*_*j,m*_ are, respectively, the scores of the query sample *i* and the reference sample *j* on the principal component *m*. The *M* eigenvalues *λ*_*m*_ are normalized to sum to one. The time complexity of the distance matrix computation is *O*(*n*_*Q*_*n*_*R*_*M*).

Given *k ≤ n*_*R*_, we denote for each query sample *i* the set of indices of its *k* nearest reference samples by 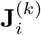. The ancestry component *q*_*i,s*_ is computed as the arithmetic mean over the ancestry components of the *k* nearest reference samples:

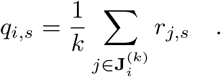

Time complexity of the ancestry estimation is dominated by sorting the distance matrix and is of the order *O* (*n*_*Q*_*n*_*R*_log(*n*_*R*_)). Therefore, the runtime of the KANN algorithm is of the order *O* (*n*_*Q*_*n*_*R*_(log(*n*_*R*_) + *M*)).

#### 2.5.2 Inverse distance weighting

In case there is a high variability among the distances between the query sample and its *k* nearest reference samples, it is plausible that the reference samples closest to the query sample are more descriptive of its ancestry than those farther away. With this in mind, we define, for a given exponent *p ≥* 0, the weight *w*(*i, j*) = *d*(*i, j*)^*−p*^ of reference sample *j* on query sample *i*, based on their inverse pairwise distance raised to the power *p*. The ancestry components are then estimated as

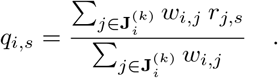

To avoid extremely large weights, KANN can be provided with a minimum distance threshold *ε* that is substituted for the distances *< ε*. In our applications, we set *ε* = 0.1.

### 2.6 Total variation distance

Ideally, for the same input data, KANN would estimate similar ancestry profiles as the current state-of-the-art haplotype-based ancestry estimation methods. To assess the performance of KANN, we compare our estimated sample profiles with the SOURCEFIND profiles for the same samples. As a distance measure between two *S*-dimensional ancestry profiles 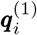 and 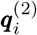 we use the total variation distance (TVD), defined as

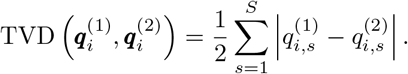

### 2.7 Model building and test data sets

From the 18,125 samples, we extracted a test set of 1,000 samples that did not overlap with the 1,798 samples used to represent the Finnish source populations in the SOURCEFIND analysis. To ensure that all 10 source populations were represented in the test set, for each population, we randomly selected 5 such samples that had *≥* 75% of their ancestry assigned to the population by SOURCEFIND. This property of the test set holds also with respect to either two or six source populations. The remaining 950 samples in the test set were selected randomly. The 17,125 samples not included in the test set are referred to as the model building data set.

### 2.8 Parameter optimization

We optimize parameters *k* and *p* within the model building data set by minimizing TVD between our estimated ancestry profiles and those available for the same samples from SOURCEFIND. Hereafter, unless otherwise specified, we use “TVD” to refer to the total variation distance between the SOURCEFIND profile and an ancestry profile estimated using some other method.

This optimization process is repeated separately for 2, 6 and 10 source populations, and for three different ways to define the reference profiles. In each setting, the model building data set is divided into reference samples and query samples. Instead of an exhaustive search through all possible parameter values, we apply the optimization in a restricted parameter space. For the parameter *p*, we ran KANN for integer values from 0 to 5. When *p* = 0, an equal weight is assigned to all pairwise distances, corresponding to the case where no distance weighting is applied. Depending on the setting, we ran the algorithm with different increments of *k* starting from 1. The three scenarios for defining the reference profiles are as follows.

#### DISCRETE

In the DISCRETE scenario, the reference samples are those with a discrete assignment to a single population, i.e., 100% of the ancestry of a reference sample is assigned to one source population. This approach enables a meaningful comparison between KANN and such ancestry inference tools that require discrete reference profiles. In our data, we have discrete population assignments for the samples that were used as the reference samples in the original SOURCEFIND analysis. For parameter *k*, we examine values 1, 10, 25, 50, 100, 250, 500, 1,000, and *n*_*R*_, where *n*_*R*_ is the maximum number of reference samples available in each set. This equals to 1,443 samples (2 populations), 1,006 samples (6 populations) and 1,214 samples (10 populations).

#### CONTINUOUS

In the CONTINUOUS scenario, we use the same reference set as in the DISCRETE scenario, namely 1,443 samples (2 populations), 1,006 samples (6 populations) and 1,214 samples (10 populations), and the same range of values for the parameter *k*. The difference here is that we utilize the continuous ancestry profiles estimated by SOURCEFIND, where the individual’s profile consists of proportions of ancestry assigned to all *S* source populations. This way we can demonstrate KANN’s ability to utilize reference samples with continuous ancestry profiles.

#### CONTINUOUSALL

The CONTINUOUSALL scenario is an extension of the CONTINUOUS scenario, where we again utilize information on continuous ancestry profiles, but use all available samples from the model building data set as our reference set. For parameter optimization, we use cross-validation. First, the samples are randomly assigned to 10 cross-validation folds, each containing approximately 10% of the samples. Each fold in turn is considered as the query samples, and the remaining folds as the reference samples. Compared to the DISCRETE and CONTINUOUS scenarios, we now have a larger number of reference samples available. We extend the parameter space for *k* to cover values 1, 3, 5, 10, 25, 50, 100, 250, 1,000, 5,000, 10,000 and 15,000.

The optimal pair of parameters in the DISCRETE and CONTINUOUS scenarios is the one that minimizes the mean TVD among the query samples, and in the CONTINUOUSALL scenario, the one that minimizes the mean TVD over the combined set of query profiles from the 10 cross-validation folds. After the parameter optimization, we estimate ancestries for the test set of 1,000 samples using KANN with different numbers of source populations and for the three scenarios of reference profiles. In each case, the algorithm is run with the parameter pair found during the optimization process.

### 2.9 Comparison with Rye

The accuracy and runtime of KANN was compared to Rye, a recent software tool for genetic ancestry inference based on principal components applicable at a biobank scale [9]. Rye uses Markov chain Monte Carlo (MCMC) optimization to find vectors in the PC-space to represent each of the ancestral source populations. Genetic ancestry profiles for the query samples are then estimated by regressing their PC vectors on those of the source populations using non-negative least squares regression (NNLS). Rye was shown to be comparable in accuracy with RFmix [6], ADMIXTURE [8] and iAdmix [7], and outperforming them with respect to runtime [9].

We used Rye to estimate ancestry profiles for the 1,000 test set samples with respect to 2, 6 and 10 Finnish source populations, and compared the estimated profiles with SOURCEFIND using TVD. In all cases, we used 10 PCs and the default parameters. Rye, like many ancestry inference methods, uses reference samples allocated to a single source population. Thus, we used the same reference sample set for Rye as we used in our DISCRETE scenario, because those are the only samples in our data with a discrete assignment to source populations. This enables a meaningful comparison between Rye and KANN, since identical input data is used with both methods.

Further, we compared the runtime between Rye and KANN in estimating the test set profiles with respect to 10 source populations. Both methods used the same 1,214 reference samples with discrete ancestry information. KANN’s runtime includes the pairwise distance matrix computation between the reference and test samples. KANN was run with parameters *k* = *n*_*R*_ *−* 1 = 1, 213 and *p* = 3. (As KANN is exceptionally fast in the special case *k* = *n*_*R*_, where the nearest neighbor search can be skipped, for this runtime comparison, we ran KANN using *k* = *n*_*R*_ *−* 1 instead of the optimized value *k* = *n*_*R*_.) Taking into account the possible variation in runtimes, we report the median and range of runtimes from 10 separate runs for both algorithms. Runtime was measured using an Intel Xeon Gold 6148 CPU and 173GB of RAM, with KANN first running on a single core and then on four cores, while Rye utilized up to four cores.

### 2.10 Interpreting the TVD values

TVD between individual’s profile estimated by KANN and SOURCEFIND tells which proportion of ancestry KANN has assigned to different source populations than SOURCEFIND. Thus, small TVD values between KANN and SOURCEFIND indicate that KANN can provide similar accuracy to a state-of-the-art haplotype-based ancestry estimation method. As TVD adds up the absolute differences over all ancestry components, it does not directly tell for which population pairs the difference occurred. We studied further the magnitude of the differences for individual source populations, and determined whether some pairs of the source populations get mixed up by the two methods more often than others. These analyses were conducted on the test set where the ancestry profiles were estimated using the parameter values optimized in the model building phase.

First, we looked at the marginal differences KANN *−* SOURCEFIND between the individual ancestry components in the test set. Furthermore, we studied how the marginal differences of the other populations are distributed among the samples that have a high marginal difference in one population. In turn, we took each source population as our target population, and identified the test set samples with a marginal difference 0. *≥* 05 for that population. The individuals with a high marginal difference in the target population are divided into two groups based on the sign of the difference (positive or negative). A positive difference implies that, compared to SOURCEFIND, KANN has overestimated the ancestry component of the population. Correspondingly, a negative difference implies that KANN has underestimated the component. For the samples in each group, we calculated the arithmetic means of the marginal differences with respect to all other populations. We were interested which populations had marginal difference to the opposite directions than our target population, as that suggests that those populations and our target population had got mixed up between KANN and SOURCEFIND. We normalized the absolute values of the mean differences that were to the opposite direction compared to the target population’s difference, and we ignored the possible differences with the same sign as the difference of the target population.

## 3 Results

### 3.1 Optimal parameters

The parameter optimization yields a pair of parameters (*k, p*) that minimizes the mean TVD in the query data set (Table 1). The smallest optimal *p* encountered was 3, indicating that the ancestry estimation benefits from the distance weighting. In the DISCRETE and CONTINUOUS scenarios, the optimal number of nearest reference samples *k* was always relatively close to *n*_*R*_, the total number of the reference samples. When a larger reference sample set was used in the CONTINUOUSALL scenario, the optimal *k* was much smaller (*k* = 25 or *k* = 50).

**Table 1:** Optimal values of *p* and *k* for Rye and the three versions of KANN for different numbers of source populations *S*. Mean TVD on the test set with 95% confidence interval is reported.

| **RYE** |  |  |  |
| --- | --- | --- | --- |
| $S$ | **Mean TVD** | | |
| 2 | 0.082 (0.077, 0.086) |  |  |
| 6 | 0.177 (0.170, 0.184) |  |  |
| 10 | 0.244 (0.237, 0.252) |  |  |

| **KANN-DISCRETE** |  |  |  |
| --- | --- | --- | --- |
| $S$ | $p$ | $k$ | **Mean TVD** |
| 2 | 3 | 1443 | 0.074 (0.07, 0.079) |
| 6 | 3 | 1000 | 0.138 (0.129, 0.146) |
| 10 | 3 | 1214 | 0.186 (0.175, 0.198) |

| **KANN-CONTINUOUS** |  |  |  |
| --- | --- | --- | --- |
| $S$ | $p$ | $k$ | **Mean TVD** |
| 2 | 4 | 1000 | 0.061 (0.058, 0.065) |
| 6 | 5 | 1000 | 0.119 (0.111, 0.126) |
| 10 | 5 | 1214 | 0.148 (0.139, 0.157) |

| **KANN-CONTINUOUSALL** |  |  |  |
| --- | --- | --- | --- |
| $S$ | $p$ | $k$ | **Mean TVD** |
| 2 | 4 | 50 | 0.036 (0.034, 0.038) |
| 6 | 3 | 25 | 0.069 (0.065, 0.074) |
| 10 | 3 | 25 | 0.112 (0.105, 0.119) |

Supplementary Fig. S4 shows the mean TVD observed across the parameter values. Regardless of the scenario, the worst results are observed when the inverse distance weighting is not applied and *k* equals to its maximum possible value *n*_*R*_. This is not surprising, as such a choice corresponds to estimating the same profile for all query samples. The weighting parameter *p* has less impact on the mean TVD when *k* is small. For example, in the KANN-CONTINUOUSALL scenario, where the optimal *k ≤* 50, changes in *p* shows less than a 0.001 difference in the mean TVD. However, in the same scenario, with a large *k* = 15, 000, we see up to a 0.35 difference in the mean TVD between values *p* = 0 and *p* = 5.

### 3.2 Ancestry estimation accuracy and runtime

Fig. 2 shows the TVD distributions of the test set profiles estimated using KANN and Rye with respect to 10 Finnish source populations. (Supplementary Fig. S5 and Supplementary Fig. S6 show the distributions with respect to 2 and 6 populations, respectively.) The box plots show a similar trend with respect to the median TVD, as Table 1 shows with respect to the mean TVD. As expected, TVD increases with the number of source populations for every method. In all cases, Rye shows the largest TVD. This is followed by KANN-DISCRETE that provides a meaningful comparison with Rye, since there KANN is run with an identical reference sample set and the same discrete ancestry information as Rye. Incorporating continuous ancestry information with KANN-CONTINUOUS decreases TVD compared to using the discrete information. Finally, extending the reference sample set to cover the whole model building data set with KANN-CONTINUOUSALL shows the smallest TVD.

**Figure 2:**
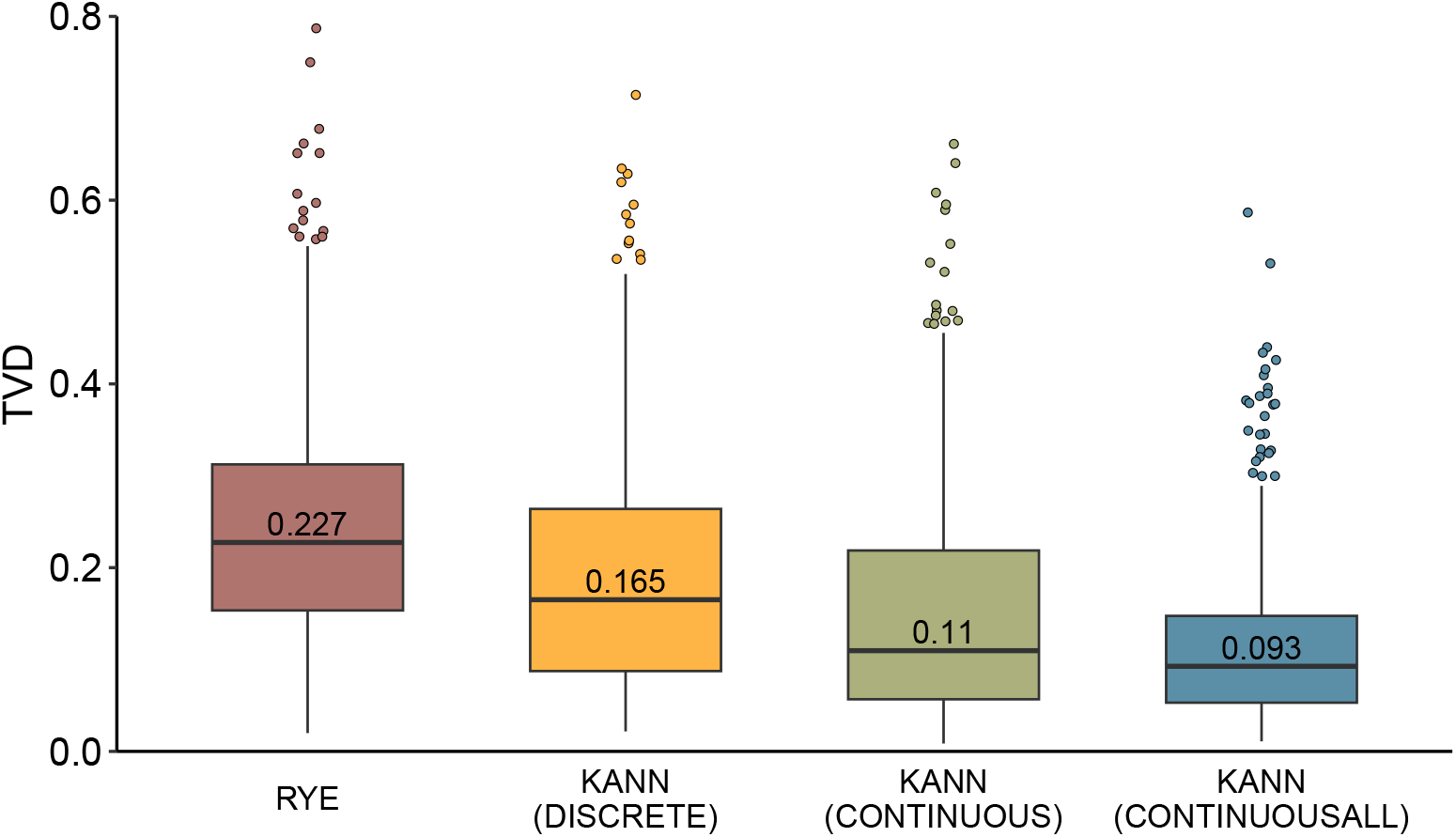
Box plots showing the TVD distribution of the test sample profiles with respect to 10 populations. Profiles are estimated using Rye, and the three versions of KANN. The profiles in each version of KANN have been estimated using the optimal parameter pair from Table 1. The median TVD is shown on each box plot.

In the test set, KANN and SOURCEFIND ancestry estimates for each individual source population had a correlation *ρ ≥* 0.859 with all 10 populations (Supplementary Fig. S7), *ρ ≥* 0.963 with 6 populations (Supplementary Fig. S8), and *ρ* = 0.988 with 2 populations (Supplementary Fig. S9). Here, the KANN profiles had been estimated using ancestry information and reference samples from the KANN-CONTINUOUSALL scenario, and the optimal parameters corresponding to the number of source populations from Table 1.

For the runtime comparison, Rye and KANN were run on an identical set of 1,214 reference samples and the 1,000 test set samples as query individuals. The median runtime for KANN over 10 runs was 9.6 seconds (min = 9.4s, max = 9.9s) using one core and decreased to 4.0 seconds (min = 3.9s, max = 4.7s) using four cores. Rye took considerably longer with a median runtime of 616 seconds (min = 541s, max = 679s) using four cores. We confirmed that the runtime of KANN was linear in the number of query samples both with one and with four cores.

### 3.3 Differences between KANN and SOURCEFIND

Population specific marginal differences between KANN and SOURCEFIND with respect to the 10 source populations are depicted in Fig. 3. These are calculated from the test sample profiles using continuous ancestry information with the whole model building dataset of 17,125 samples as the reference, and using the optimized parameters (*k* = 25, *p* = 3). For all populations, the absolute values of the lower and upper quartiles remain *≤* 0.04. East Lapland (ELA), although having a small interquartile range, shows the largest positive (0.507) and negative (-0.531) differences. Supplementary Fig. S10 and Supplementary Fig. S11 show the marginal differences with respect to 2 and 6 populations, respectively.

**Figure 3:**
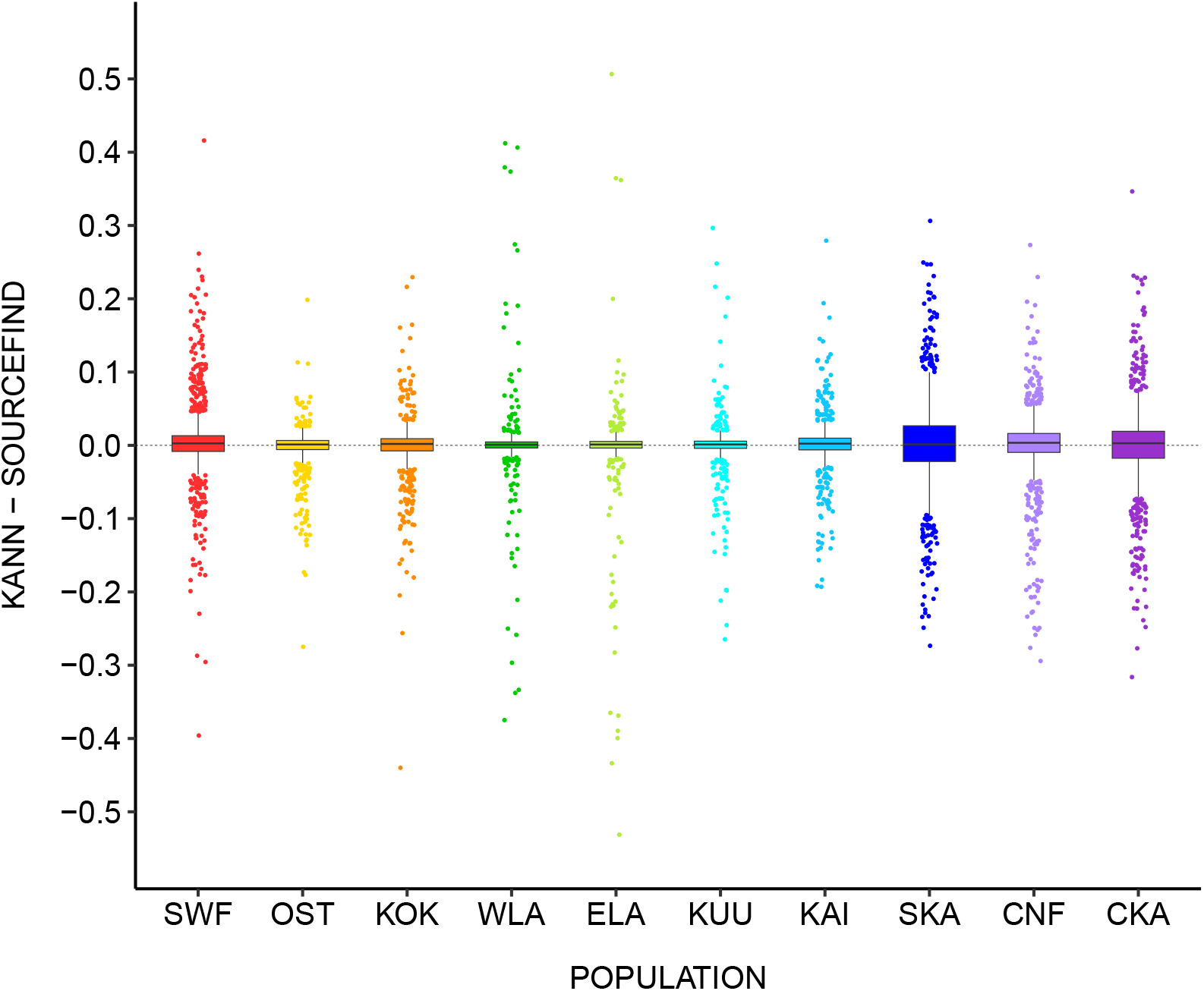
Marginal differences between KANN and SOURCEFIND in the ancestry components of the test set with respect to 10 source populations. The population abbreviations and colors are as in Fig. 1.

Using the same estimated ancestry profiles, Fig. 4 visualizes the distributions of population-specific mean marginal differences among the samples having a high marginal difference in a particular target population (Supplementary Table S1). Supplementary Fig. S12 depicts the distributions with respect to 6 populations.

**Figure 4:**
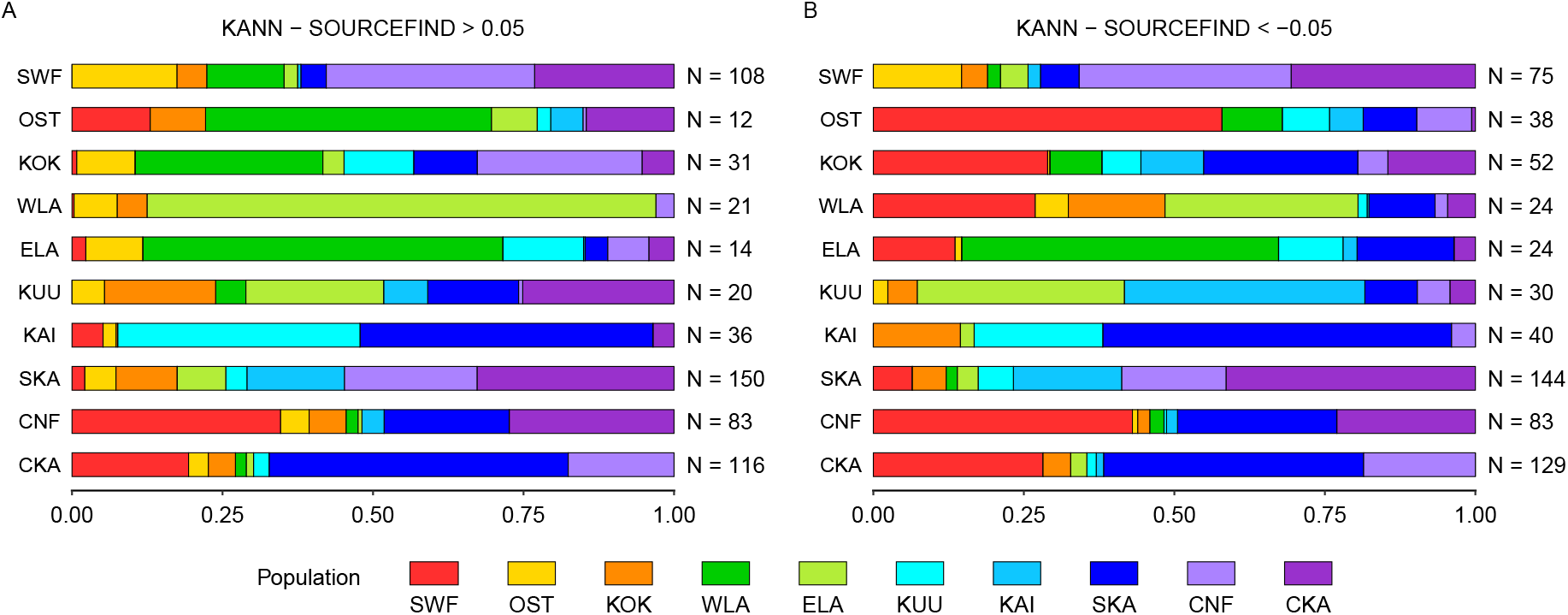
Distributions of the normalized absolute mean differences (x-axis) among the samples having a high A) positive or a high B) negative marginal difference in a particular target population. The y-axis shows the target population label (left-hand side), and the corresponding number of samples (right-hand side) reaching the threshold of 0.05 difference with respect to the target population. The population abbreviations and colors are given in Fig. 1.

In general, population pairs that are geographical neighbors (Fig. 1), are most prone to get mixed up between KANN and SOURCEFIND. Fig. 4 also indicates that the distributions in panels A and B are not entirely symmetrical. For example, among the individuals with a high positive difference in the West Lapland (WLA) component (row 4 in Fig. 4A), the vast majority (84.5%) of the ancestry was mixed up with its geographical neighbor, East Lapland (ELA), with virtually no mix-up (0.3%) with the geographically more distant Southwestern Finland (SWF). On the other hand, among the individuals with a high negative difference in the WLA component (row 4 in Fig. 4B), the ancestry has been mixed up substantially less with ELA (32.1%), while almost a comparable fraction (26.9%) of ancestry was estimated to the SWF component.

## 4 Discussion

We developed KANN with the aim to offer a user-friendly and scalable tool that could produce similar results as computationally more complex haplotype-based ancestry estimation method SOURCEFIND [5]. Our results indicate that KANN is able to estimate ancestry profiles that are well in line with SOURCEFIND. For example, with up to 6 source populations, all ancestry components from KANN and SOURCEFIND are highly correlated (*>* 0.96).

Naturally, the average difference between KANN and SOURCEFIND increases as we attempt to assign the Finnish ancestry into 10 fine-scale source populations. For many target populations, the difference between KANN and SOURCEFIND is nearly independent of which one gives a higher ancestry estimate for these target populations as indicated by the similarity of the panels A and B of Fig. 4. In these cases, the largest differences between KANN and SOURCEFIND tend to occur between source populations that are genetically close to each other as measured by the Fst values reported by Kerminen et al. in their Table S2 [13]. Instead, for three geographically western populations OST, KOK and WLA, there is a clear asymmetry between the panels A and B of Fig. 4. Compared to SOURCEFIND, KANN tends to replace some ancestry in these three populations by the SWF component (Fig. 4B) that represents the canonical Western Finland component, whereas for individuals for whom KANN’s estimate in these three populations is larger than the SOURCEFIND’s estimate, the extra ancestry is mainly taken from one of the neighboring populations but not from SWF (Fig. 4A). Such asymmetry could reflect both complex relationships between the western source populations and properties of the PCA used in the KANN algorithm that may emphasize particular aspects of the population structure of the western Finland.

KANN was able to reproduce the SOURCEFIND’s estimates more accurately than an existing PCA based ancestry estimation method Rye [9] when using the same input data. Furthermore, we achieved a considerable improvement with KANN when we extended KANN to utilize continuous ancestry profiles of the reference individuals instead of assigning each reference individual to a single source population as required by Rye. Given that individuals from natural populations are often complex mixtures of different ancestries, we consider KANN’s ability to use the continuous ancestry profiles as an important advancement for many ancestry estimation applications.

A practical question is how to choose suitable values for the two parameters required by KANN: *k* the number of nearest neighbors and *p* the exponent of the inverse distance that is used to weight the *k* nearest samples. If high quality ancestry information can be estimated for a subset of the data, for example by SOURCEFIND or some other haplotype-based method, the parameters can be optimized by using a cross-validation approach as we have demonstrated. If such information is not available, the parameters can alternatively be chosen based on some general trends we observed about the parameters effect on the estimation accuracy (Supplementary Fig. S4). The choice of *k* and *p* should be considered jointly, as their effects on estimation accuracy depend on each other. We acknowledge that the following observations are made in our data set, and they might not generalize equally well in all other cases.

We observed that *p* had a considerable effect only when *k* was fairly large, say *>* 5% of the number of reference samples. There, increasing *p* from 0 typically first decreased the TVD to SOURCEFIND but we did not find any situation where values of *p >* 5 would had anymore decreased the TVD. Hence, we consider a plausible range for *p* to be between 0 and 5. Both the smallest and the most frequent optimal value in all of our optimization scenarios was *p* = 3 which we propose as a default value for data sets where no parameter optimization is possible.

The optimal value of *k* is influenced also by the size of the reference sample set *n*_*R*_. In our examples, when *n*_*R*_ *<* 1, 500 (KANN-DISCRETE and KANN-CONTINUOUS), large values of *k* were preferred as *k* = 1, 000 was the smallest optimal value observed. On the other hand, with the largest reference sample set with *n*_*R*_ *>* 15, 000 (KANN-CONTINUOUSALL), much smaller values (*k* = 25, *k* = 50) were optimal. However, even larger values of *k* with varying levels of distance weighting produced near optimal results, suggesting that a large reference sample set does not necessarily mean that smaller values of *k* should always be preferred. For example, the difference between the optimal mean TVD and the one obtained using, say, *k* = 10, 000 and *p* = 5 in the KANN-CONTINUOUSALL scenario is only at most 0.036 regardless of the number of populations. If no distance weighting is used (*p* = 0), then we recommend avoiding values of *k* that are close to the full reference data set size, because otherwise the ancestry estimates of all individuals would become very similar as they would be (unweighted) averages over largely overlapping sets of the reference individuals.

Obvious future application areas of KANN include population genetic studies and the individuals’ own interest in their personal genetic ancestry through direct-to-consumer services. Additionally, we expect that detailed information on individual-level genetic ancestry on biobank-scale data sets will enable new epidemiological studies to separate the effect of genetic ancestry from environmental factors. Finally, since the genetic ancestry profile does affect the performance of polygenic scores [2, 3], KANN can help us to choose most suitable genetic predictions for each individual according to the individual’s detailed ancestry profile.

## 5 Data availability

The genotype data used in this study are available from THL biobank via procedures outlined in the Finnish Biobank Act and access can be obtained for biomedical research by contacting

## 7 Supplementary data

Supplementary data are available at NAR Online.

## 8 Acknowledgements

This study utilized data from Finnish Institute for Health and Welfare (THL biobank project THLBB2019 44).

## 9 Author contributions statement

J.R.: writing (original draft), software, methodology, formal analysis. M.P.: supervision, writing (original draft), conceptualization, methodology. S.K. and A.H.: Data curation, review of the manuscript.

