## Supplementary figures for "KANN: estimation of genetic ancestry profiles by nearest neighbor regression"

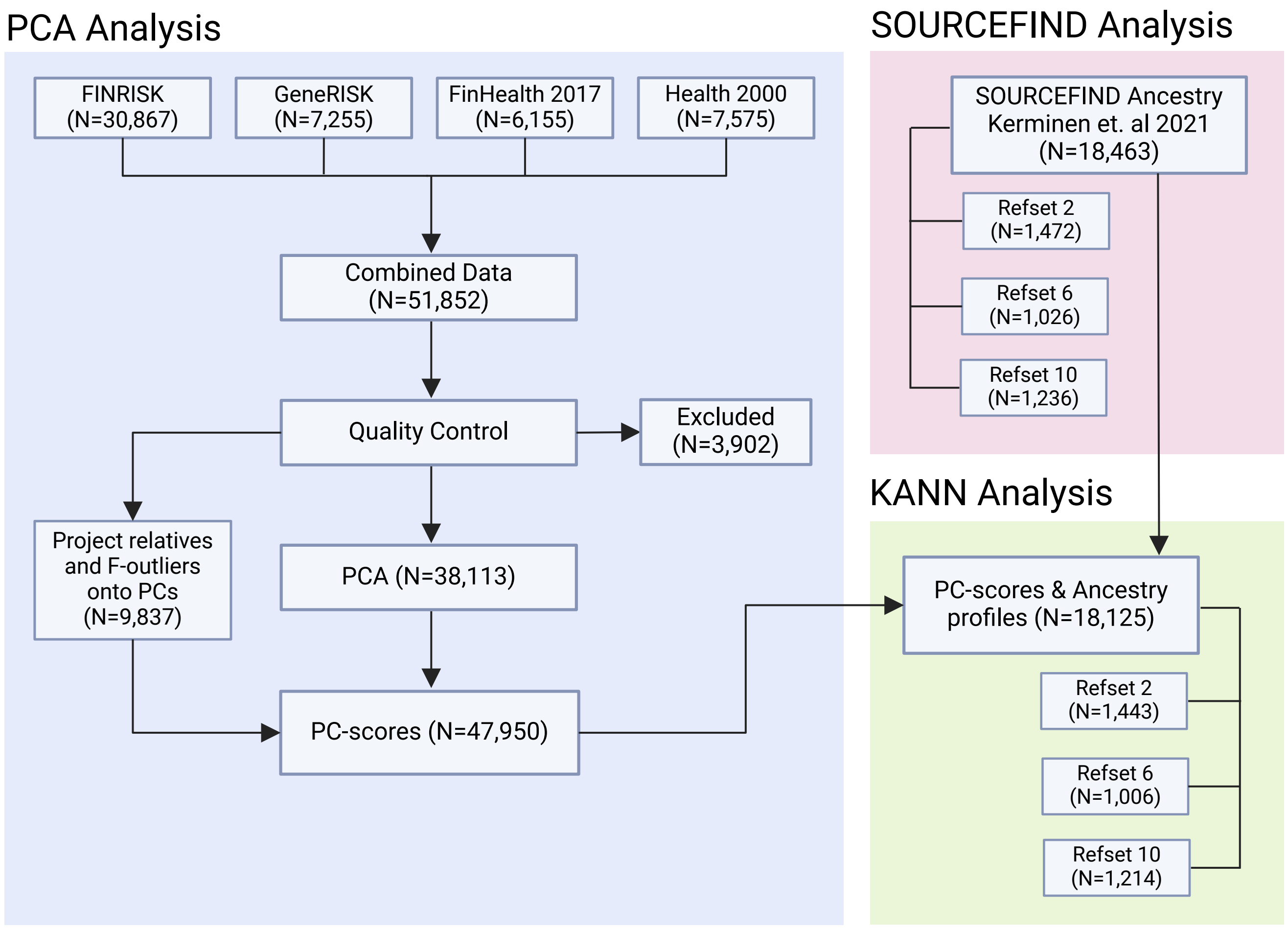

Supplementary Figure S1: Workflow of the study. Created in BioRender. Riikonen, J. (2025) https://BioRender.com/k67z392

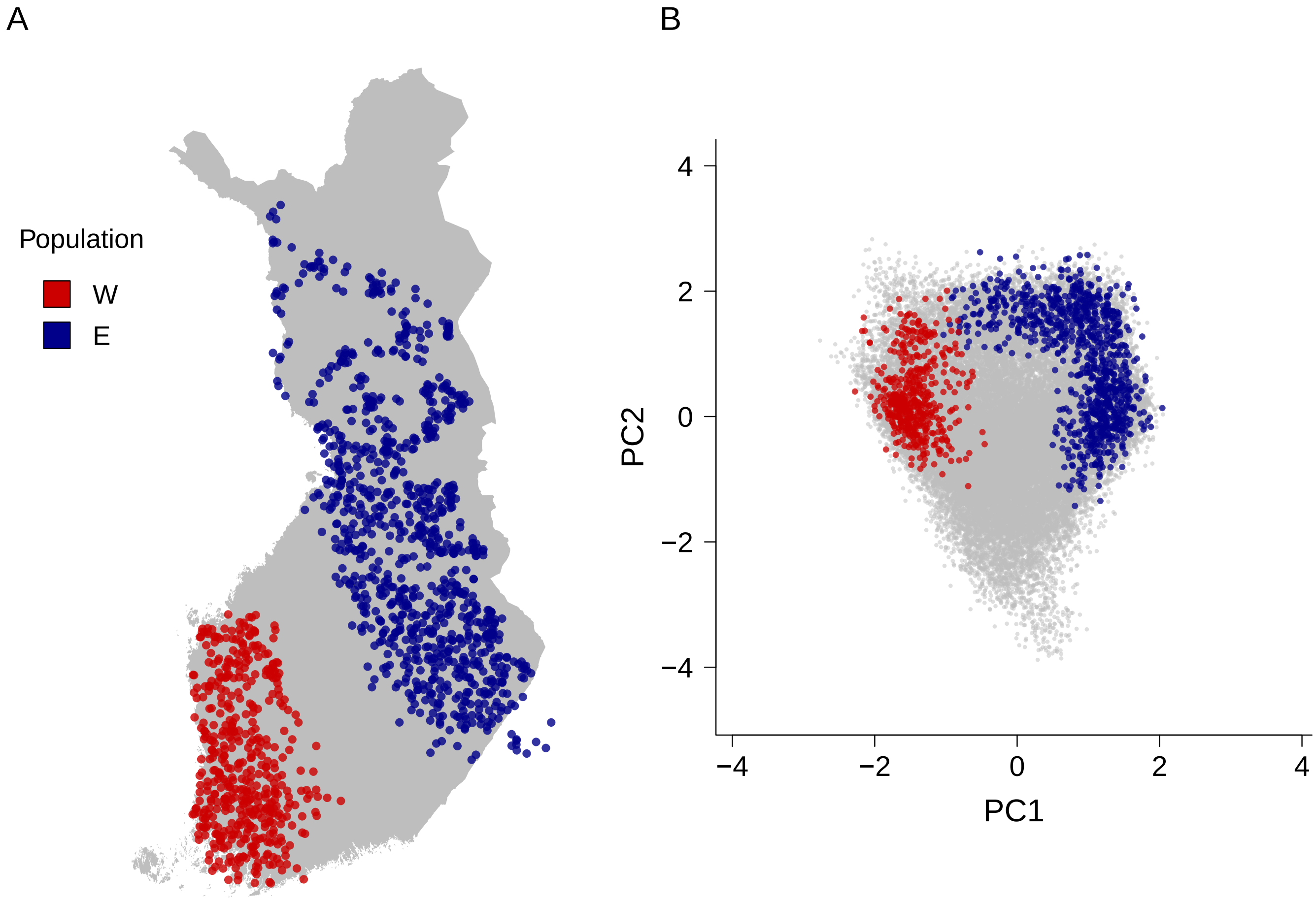

Supplementary Figure S2: A) Map of Finland with the geographical locations of the reference samples (n = 1,443) allocated to 2 Finnish source populations. The points depict the mean coordinates of the parents' municipalities of birth. B) The same individuals highlighted on the first two principal components. The samples not included in the reference groups are depicted in grey colour. W: West, E: East.

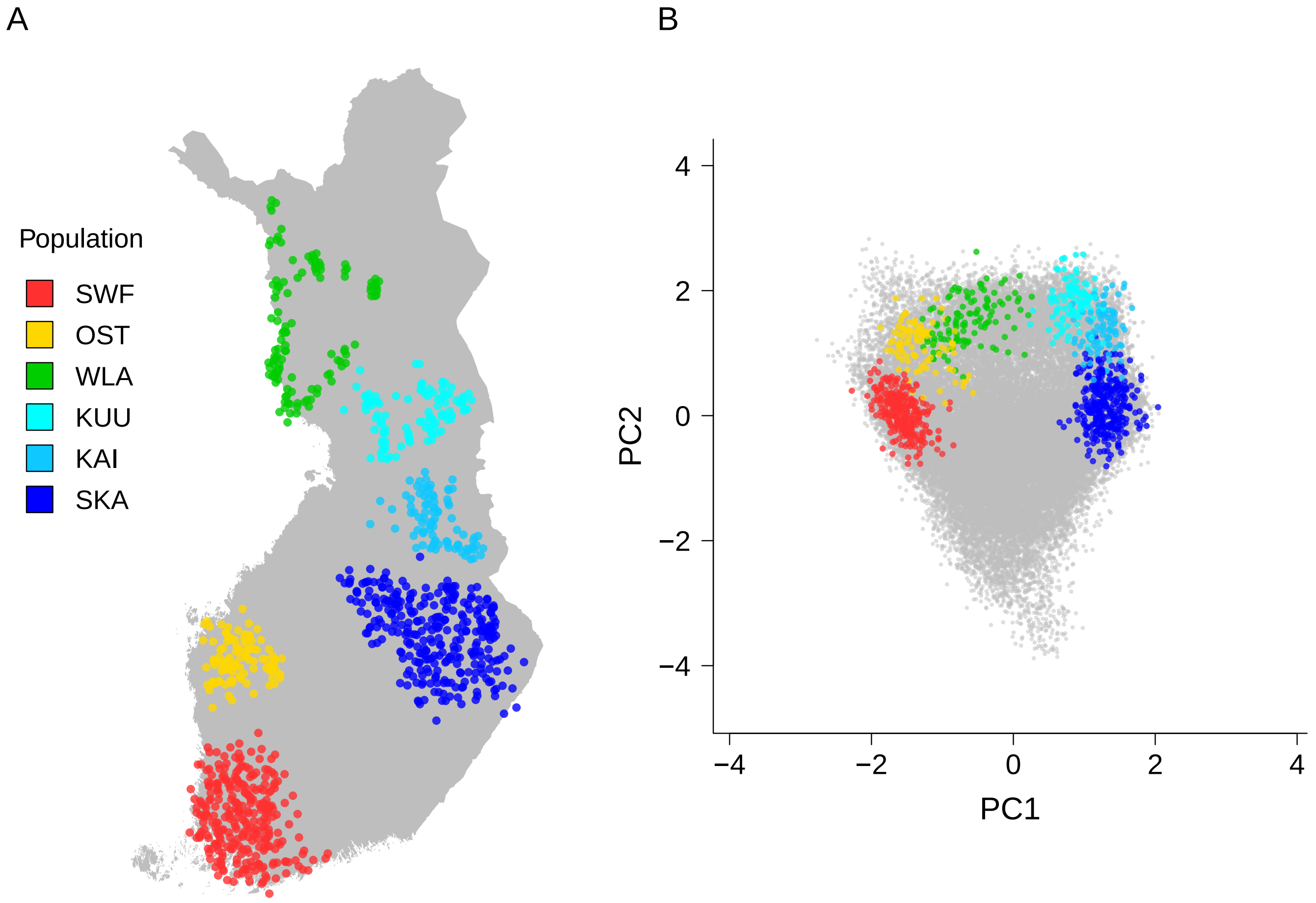

Supplementary Figure S3: A) Map of Finland with the geographical locations of the reference samples (n = 1,006) allocated to 6 Finnish source populations. The points depict the mean coordinates of the parents' municipalities of birth. B) The same individuals highlighted on the first two principal components. The samples not included in the reference groups are depicted in grey colour. SWF: Southwestern Finland, OST: Ostrobothnia, WLA: West Lapland, KUU: Kuusamo, KAI: Kainuu, SKA: Savo-Karelia.

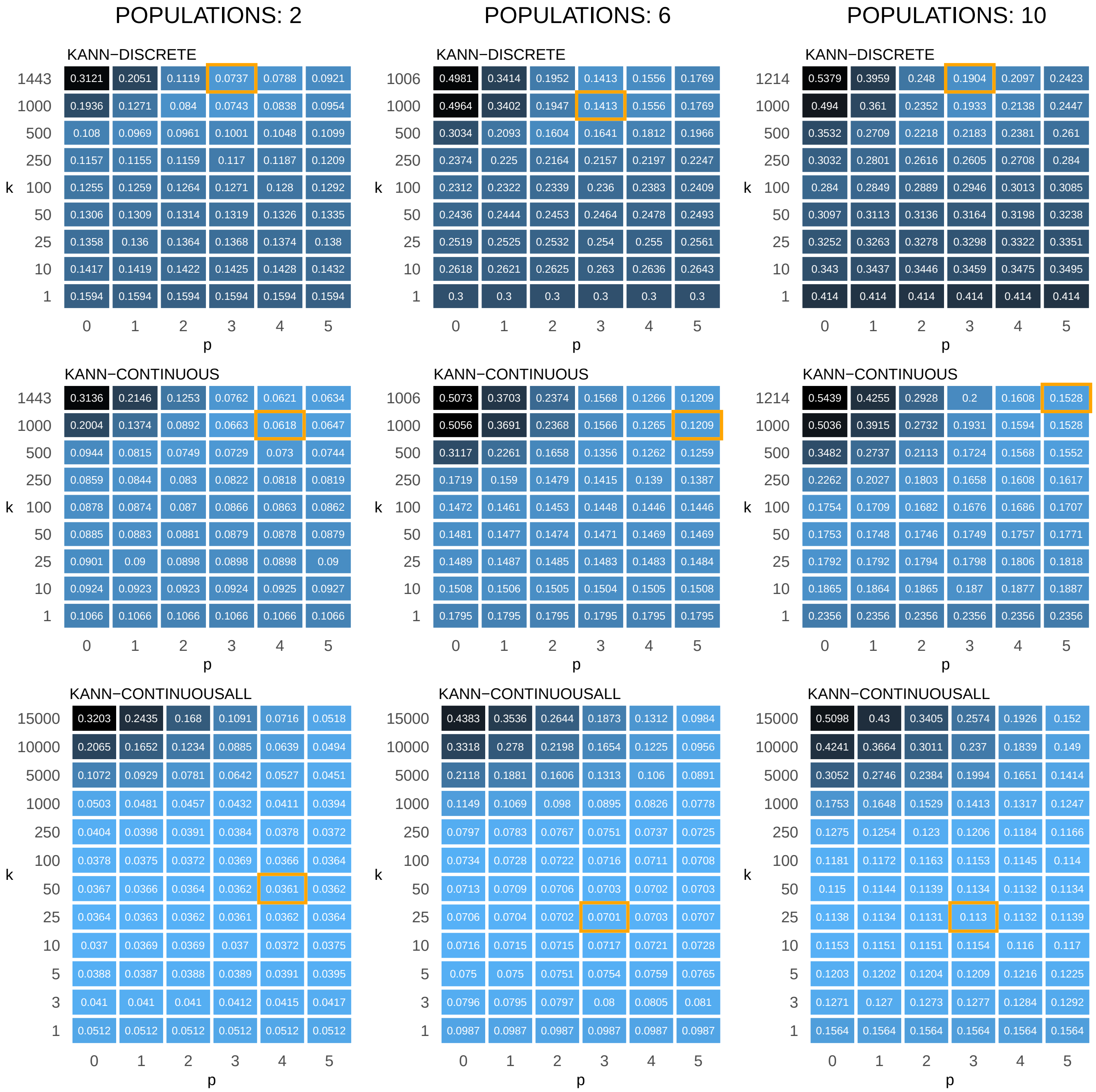

Supplementary Figure S4: Mean TVD of the query data set profiles estimated using KANN with all parameter configurations considered. The TVDs are shown for three optimization scenario (DISCRETE, CONTINUOUS, CONTINUOUSALL) and with different numbers of source populations (2, 6, 10). The values are rounded up to four decimals, with the minimum TVD highlighted.

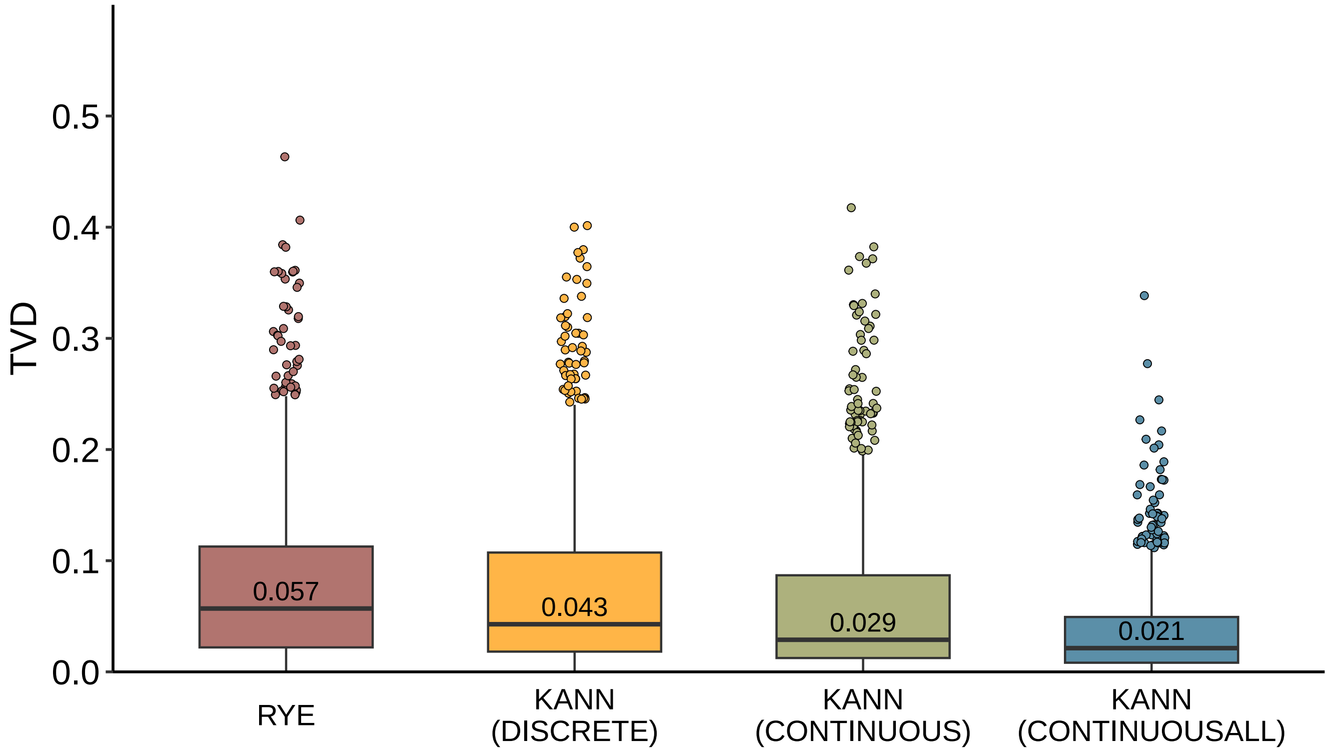

Supplementary Figure S5: Box plots showing the TVD distribution of the test sample profiles with respect to 2 populations. Profiles are estimated using Rye, and the three versions of KANN. The profiles in each version of KANN have been estimated using the optimal parameter pair. The median TVD is shown on each box plot.

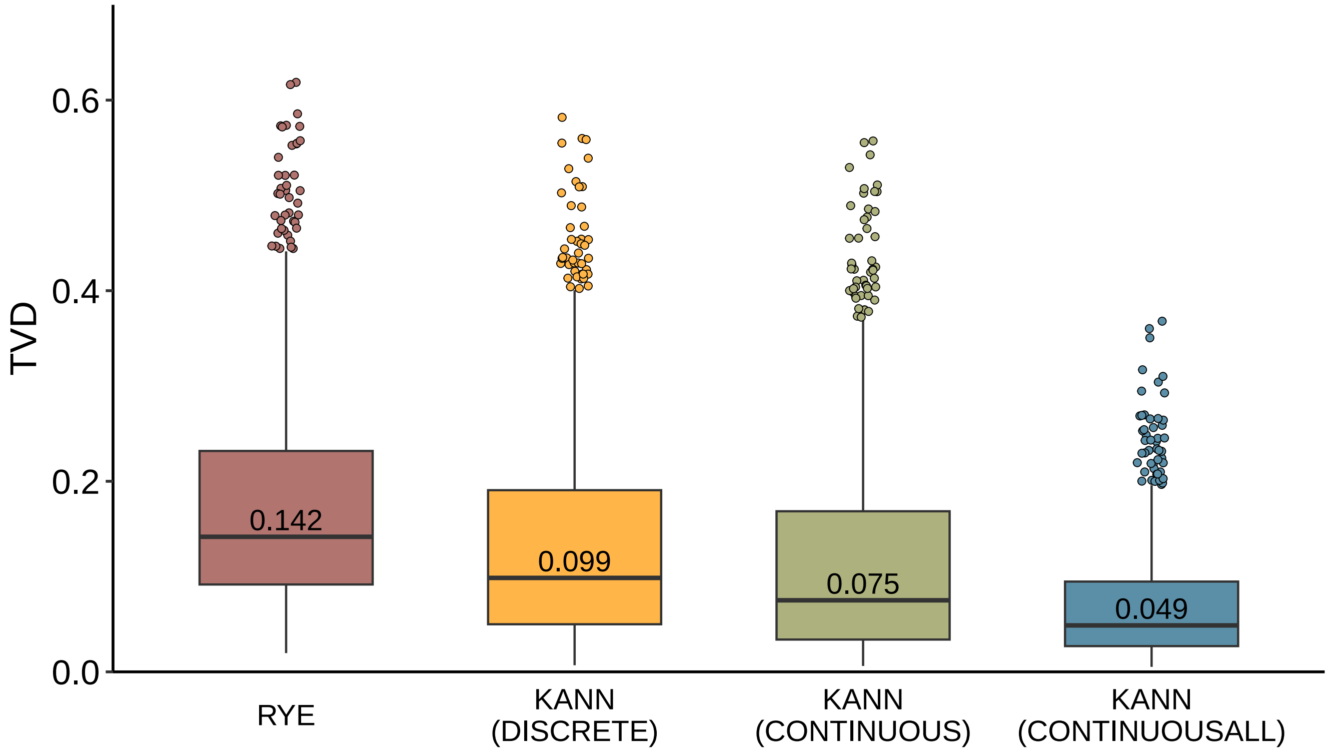

Supplementary Figure S6: Box plots showing the TVD distribution of the test sample profiles with respect to 6 populations. Profiles are estimated using Rye, and the three versions of KANN. The profiles in each version of KANN have been estimated using the optimal parameter pair. The median TVD is shown on each box plot.

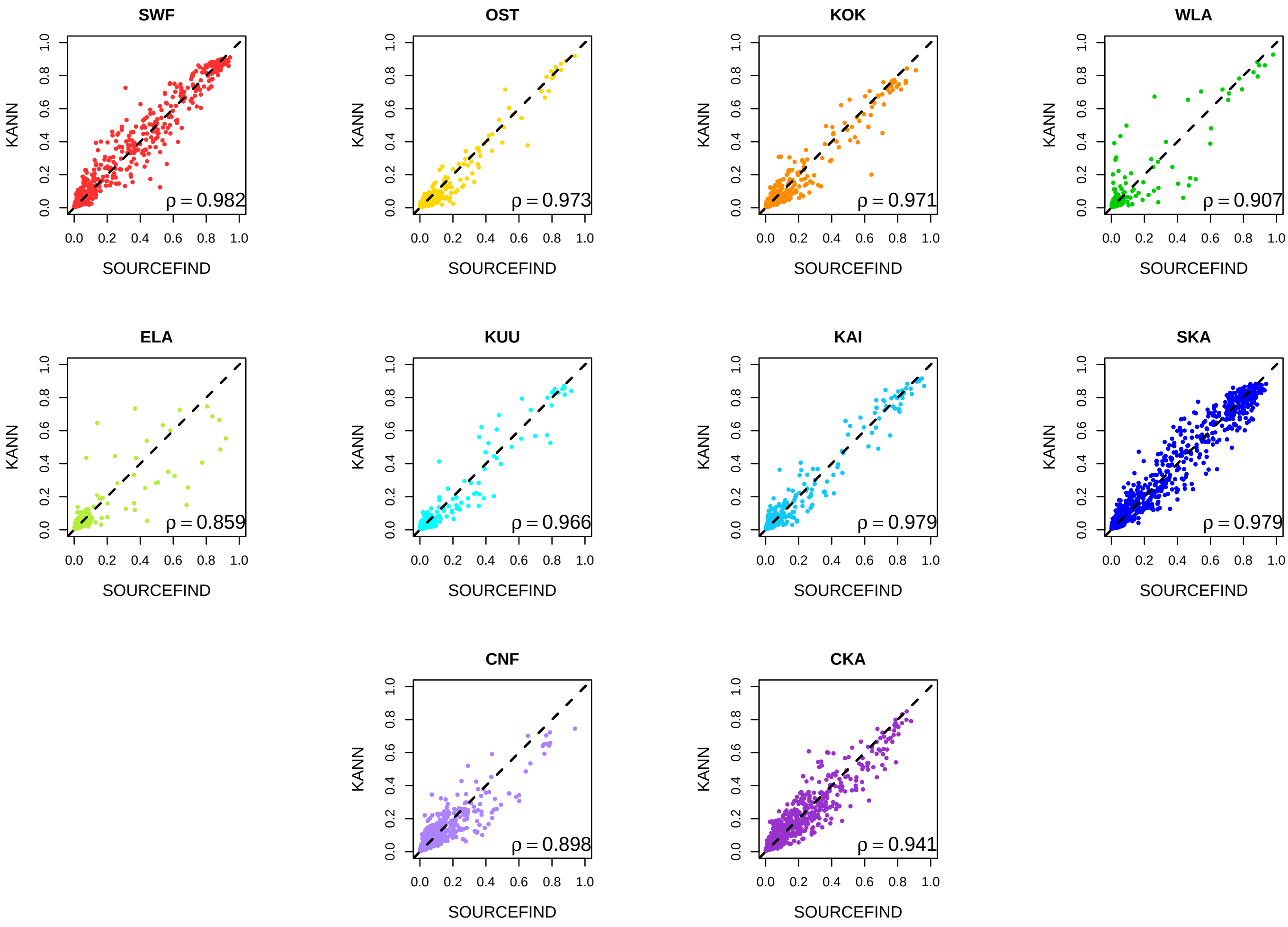

Supplementary Figure S7: Pairwise scatterplots of the test set samples' ancestry components estimated using KANN and SOURCEFIND for 10 source populations. Diagonal is shown as a dashed line. Each panel reports the Pearson correlation coefficient ($\rho$) between the estimates of the two methods. The population abbreviations and colors are given in Fig. 1.

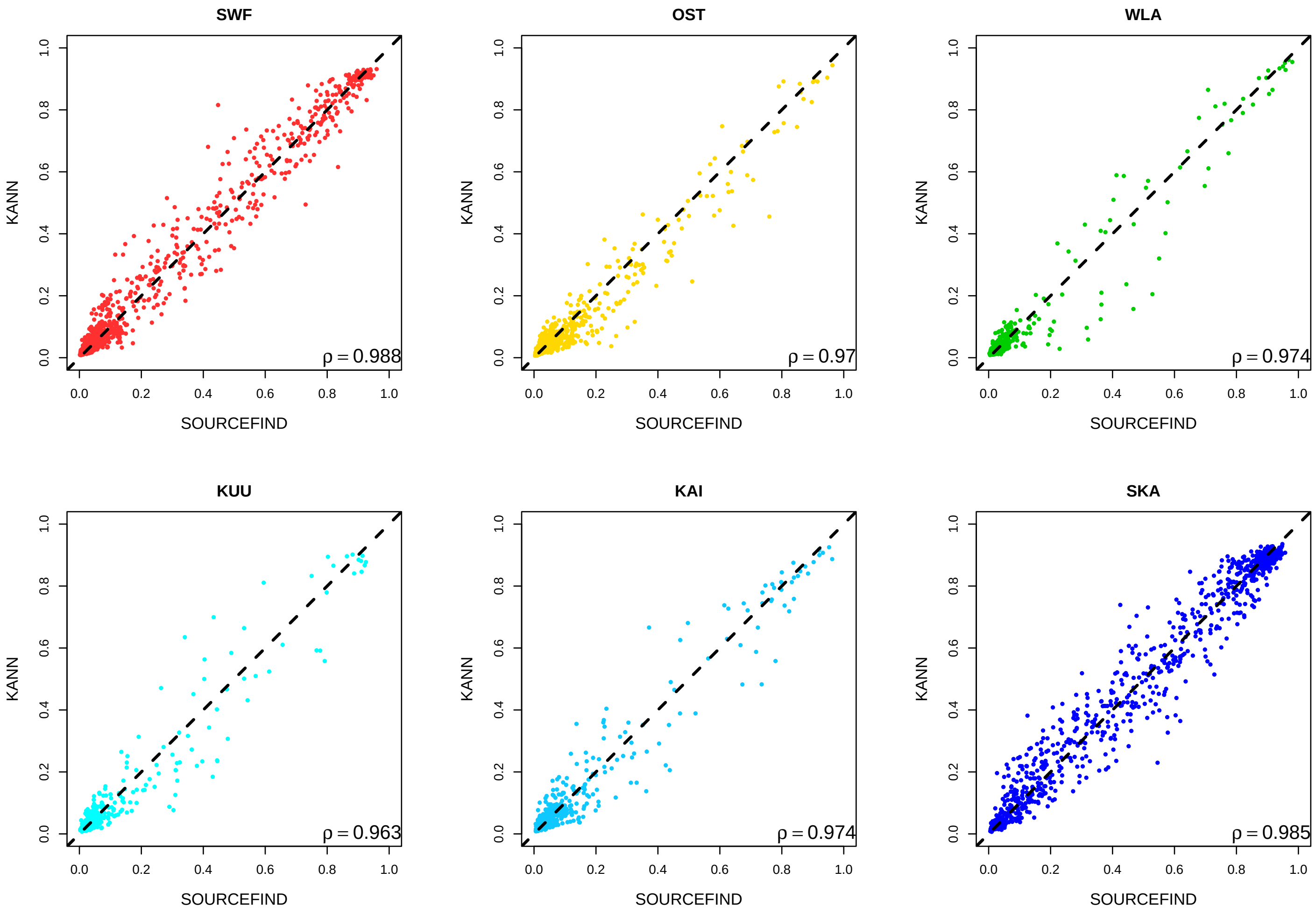

Supplementary Figure S8: Pairwise scatterplots of the test set samples' ancestry components estimated using KANN and SOURCEFIND for 6 source populations. Diagonal is shown as a dashed line. Each panel reports the Pearson correlation coefficient ($\rho$) between the estimates of the two methods. The population abbreviations and colors are given in Supplementary Fig. S3

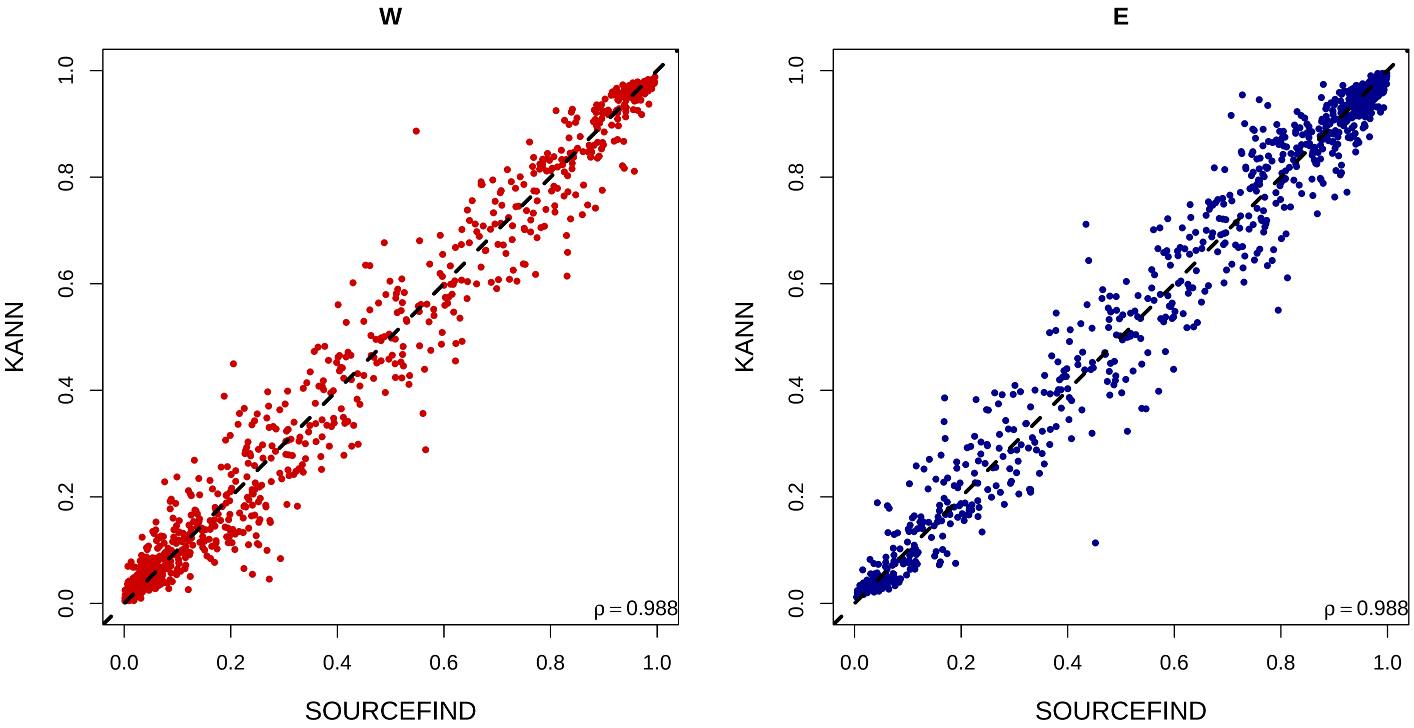

Supplementary Figure S9: Pairwise scatterplots of the test set samples' ancestry components estimated using KANN and SOURCEFIND for 2 source populations. Diagonal is shown as a dashed line. Each panel reports the Pearson correlation coefficient ($\rho$) between the estimates of the two methods. The population abbreviations and colors are given in Supplementary Fig. S2

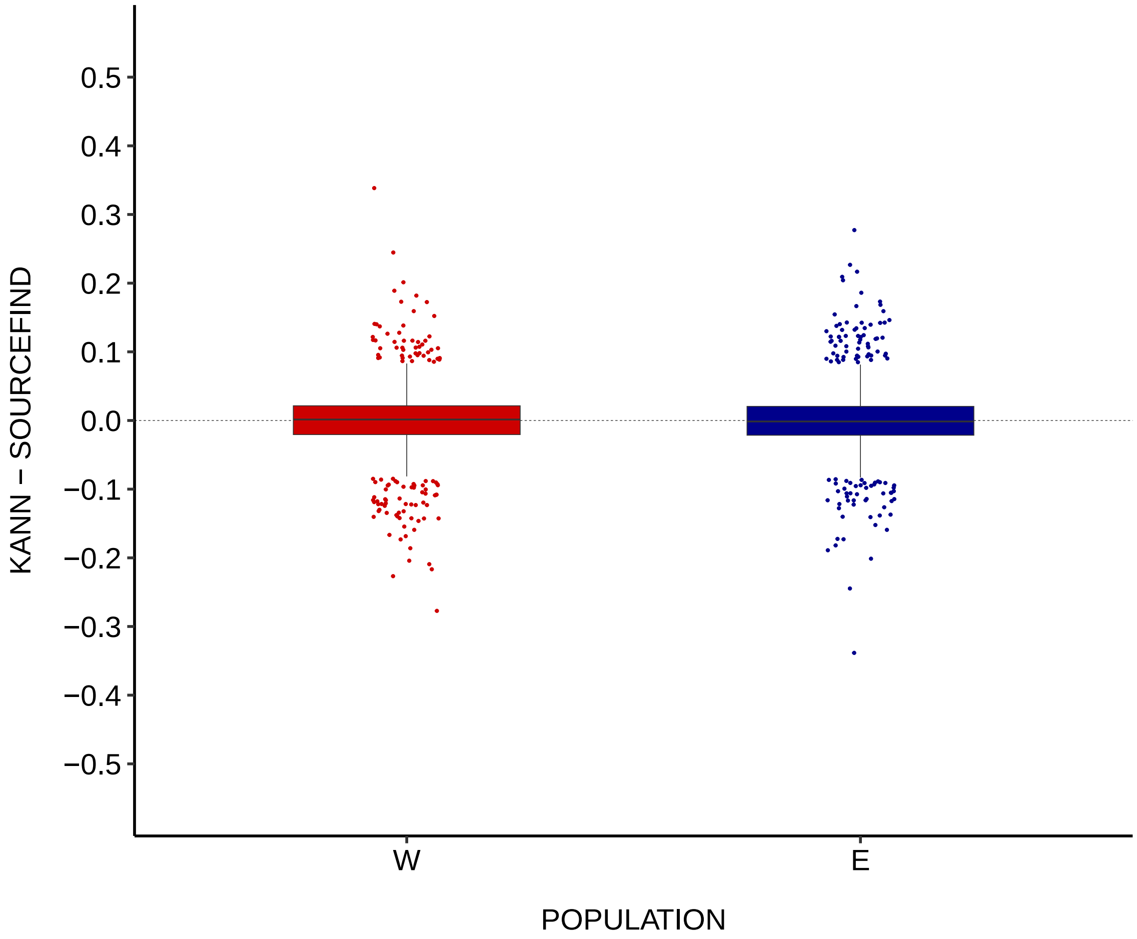

Supplementary Figure S10: Marginal differences between KANN and SOURCEFIND in the ancestry components of the test set with respect to 2 source populations. KANN profiles are estimated using 17,125 reference samples with continuous ancestry information, and the optimal parameter pair (k = 50, p = 4). The population abbreviations and colors are as in Supplementary Fig. S2.

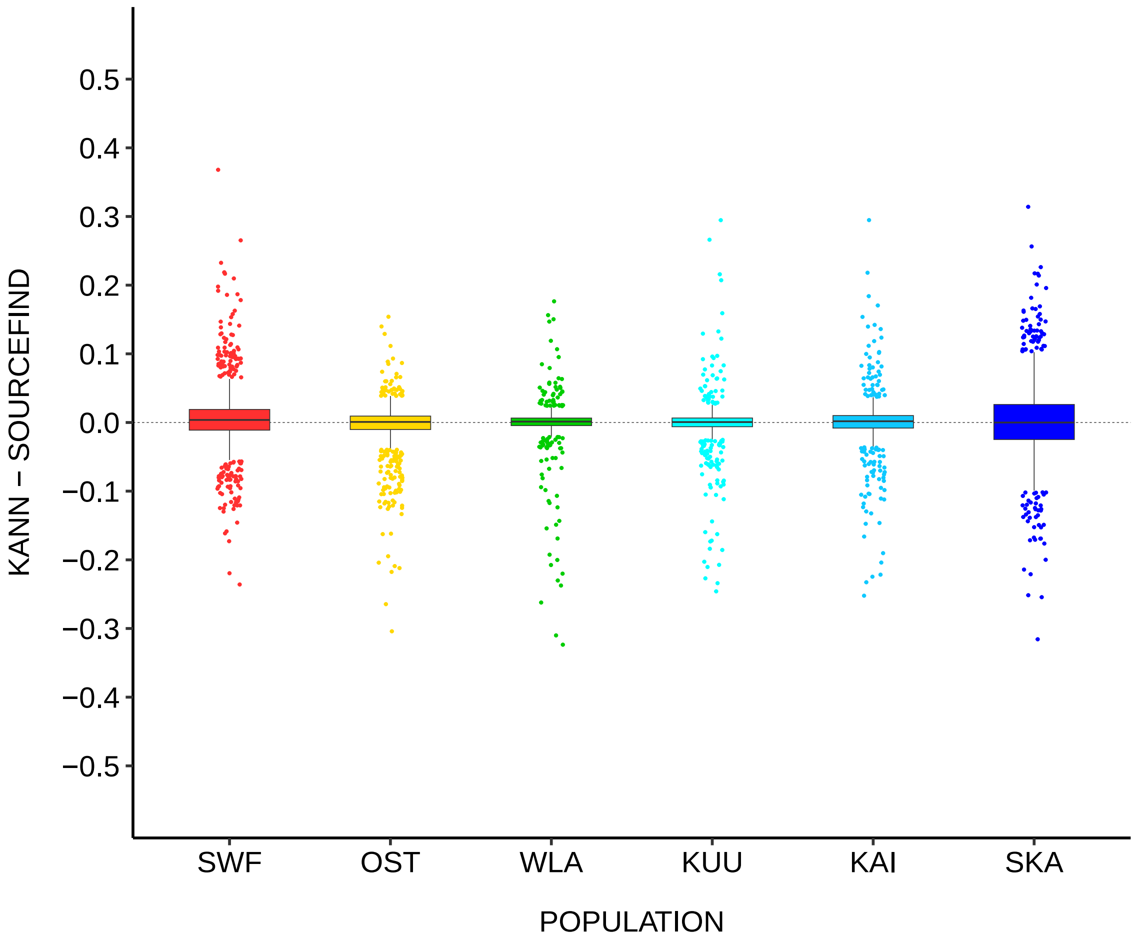

Supplementary Figure S11: Marginal differences between KANN and SOURCEFIND in the ancestry components of the test set with respect to 6 source populations. KANN profiles are estimated using 17,125 reference samples with continuous ancestry information, and the optimal parameter pair (k = 25, p = 3). The population abbreviations and colors are as in Supplementary Fig. S3.

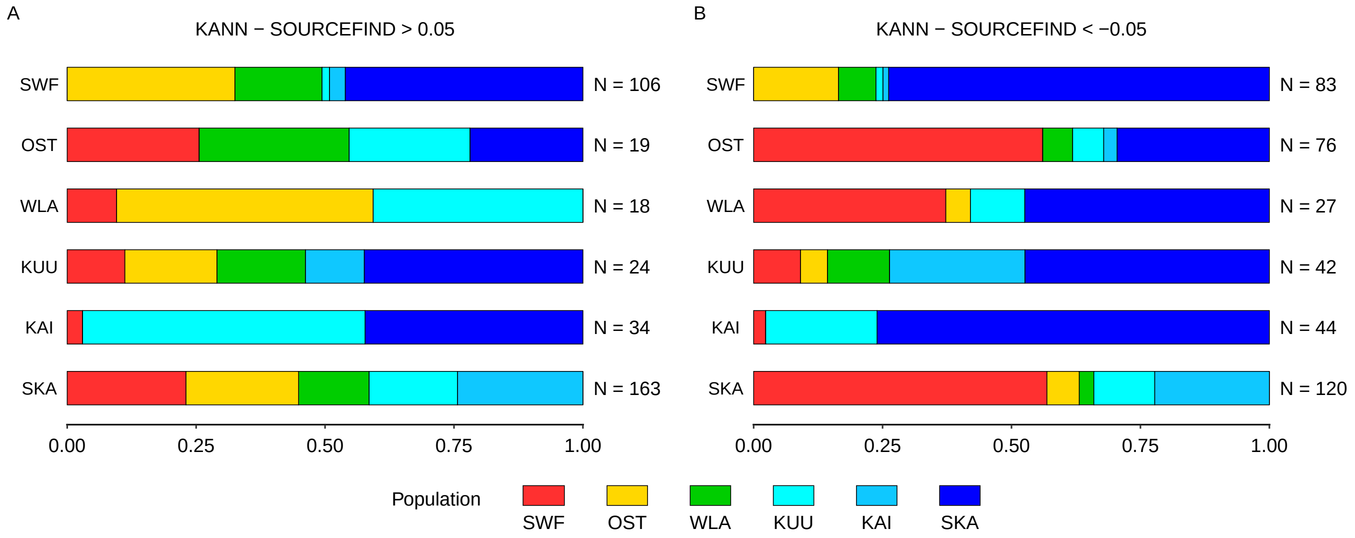

Supplementary Figure S12: Distributions of the normalized absolute mean differences (x-axis) among the samples having a high A) positive or a high B) negative marginal difference in a particular target population. The y-axis shows the target population label (left-hand side), and the corresponding number of samples (right-hand side) reaching the threshold of 0.05 difference with respect to the target population. KANN profiles are estimated using 17,125 reference samples with continuous ancestry information, and the optimal parameter pair (k = 25, p = 3). The population abbreviations and colors are given in Supplementary Fig. S3.
